# Blockade of stromal Gas6 alters cancer cell plasticity, activates NK cells and inhibits pancreatic cancer metastasis

**DOI:** 10.1101/732149

**Authors:** Lucy Ireland, Teifion Luckett, Michael C. Schmid, Ainhoa Mielgo

## Abstract

Pancreatic ductal adenocarcinoma (PDA) is one of the deadliest cancers due to its aggressive and metastatic nature. PDA is characterized by a rich tumor stroma with abundant macrophages, fibroblasts and collagen deposition that can represent up to 90% of the tumor mass. Activation of the tyrosine kinase receptor AXL and expression of its ligand growth arrest-specific protein 6 (Gas6) correlate with a poor prognosis and increased metastasis in pancreatic cancer patients. Gas6 is a multifunctional protein that can be secreted by several cell types and regulates multiple processes, including cancer cell plasticity, angiogenesis and immune cell functions. However, the role of Gas6 in pancreatic cancer metastasis has not been fully investigated. In these studies we find that, in pancreatic tumors, Gas6 is mainly produced by tumor associated macrophages (TAMs) and cancer associated fibroblasts (CAFs) and that pharmacological blockade of Gas6 partially reverses epithelial-to-mesenchymal transition (EMT) of tumor cells and supports NK cell activation, thereby inhibiting pancreatic cancer metastasis. Our data suggest that Gas6 simultaneously acts on both the tumor cells and the NK cells to support pancreatic cancer metastasis. This study supports the rationale for targeting Gas6 in pancreatic cancer and use NK cells as a potential biomarker for response to anti-Gas6 therapy.

## Introduction

Growth arrest-specific gene 6 (Gas6) is a multifunctional factor that regulates several processes in normal physiology and pathophysiology [1].Gas6 binds to the Tyro3, Axl and Mer (TAM) family of receptor tyrosine kinases (TAM receptors) with the highest affinity for Axl [2]. Gas6 supports erythropoiesis, platelet aggregation, angiogenesis, efferocytosis, and inhibits the immune response [3]. Gas6 is critical for the maintenance of immune homeostasis and mice deficient in Gas6 or TAM receptors experience severe autoimmune diseases [4]. Gas6 and its main receptor Axl are overexpressed in several cancer types including, breast, ovarian, gastric, glioblastoma, lung and pancreatic cancer and their expression correlates with a poor prognosis [5]. Axl is ubiquitously expressed in all tissues [6] but is particularly notable in cancer cells, macrophages, dendritic cells and natural killer cells for its role in driving immunosuppression and tumor progression [7–9]. Several cancer studies have focused on the role of Gas6-Axl signaling on the tumor cells and have demonstrated that Axl activation supports tumor cells proliferation, epithelial-mesenchymal transition (EMT), drug resistance, migration and metastasis [5]. Factors secreted within the tumor microenvironment are able to sustain Gas6/Axl signaling. Hypoxia Inducible Factor (HIF) has been shown to bind to the *Axl* promoter region and upregulate its expression on renal cell carcinoma cells [10]. Secretion of IL-10 and M-CSF by tumor cells induces tumor associated macrophages to secrete Gas6 [11].

However, only a few studies have investigated the role of Gas6-Axl signaling in the immune response to breast cancer, ovarian cancer and melanoma [7, 9].

In solid tumors such as breast or pancreatic cancer, the tumor stroma can represent up to 80% of the tumor mass and actively influences cancer progression, metastasis [12–14] and resistance to therapies [15–17].

Pancreatic ductal adenocarcinoma (PDA) is one of the most lethal cancers worldwide and better therapies are urgently needed [18]. Metastasis, therapy resistance, and immunosuppression are key characteristics of pancreatic tumors [19, 20]. The Gas6–Axl pathway is activated in 70% of pancreatic cancer patients [21] and is associated with a poor prognosis and increased frequency of distant metastasis [22]. Blocking Gas6 or its receptor Axl inhibits cancer progression [23, 24] and several Axl inhibitors are currently being tested in cancer patients, including PDA patients. While the cancer cell autonomous functions of Gas6 are well documented, the effect of Gas6 signaling in the stroma/immune compartment in pancreatic cancer has not been fully explored. In these studies, we sought to understand the effect of Gas6 blockade in both the tumor and the stroma/immune compartments, *in vivo*, in pancreatic cancer. Gaining a better understanding of how blockade of Gas6 signaling affects pancreatic cancer is important because it will help design and interpret the results of the recently launched clinical trials that are testing anti-Gas6/TAM receptors therapies in pancreatic cancer patients [25].

## Results

### Pharmacological blockade of Gas6 inhibits spontaneous pancreatic cancer metastasis

To investigate the effect of Gas6 blockade in pancreatic cancer growth and metastasis, we used an orthotopic syngeneic pancreatic cancer model, in which pancreatic cancer cells derived from the gold standard genetic mouse model of pancreatic cancer (LSL-Kras^G12D^; LSL-Trp53^R172H^; Pdx1-Cre mice; KPC model), transduced with a reporter lentivirus expressing zsGreen/luciferase, were orthotopically implanted into the pancreas of syngeneic immuno-competent mice. This model faithfully recapitulates features of the human disease, and tumors are highly infiltrated by macrophages and are rich in fibroblasts [16, 26, 27]. Importantly, pancreatic tumors from this mouse model also showed expression and activation of Axl receptor (Supplementary Figure 1A). These mice were then treated with isotype control IgG antibody or an anti-Gas6 neutralizing antibody (Figure 1A). 30 days after implantation, pancreatic tumors, lungs, livers and mesenteric lymph nodes were surgically removed and analysed. As expected, control treated mice showed high levels of Axl receptor activation in tumors, whereas the anti-Gas6 treated group showed markedly reduced levels of Axl receptor activation, confirming that anti-Gas6 antibody has reached the tumor and has blocked Axl signaling (Supplementary Figure 1A). No differences were seen in primary pancreatic tumor growth (Figure 1B) between the control and anti-Gas6 treatment groups. However, mice treated with the anti-Gas6 antibody showed reduced metastasis to lungs, livers and mesenteric lymph nodes, compared to control treated mice, as assessed by biolumiscence *ex-vivo* imaging of these organs (Supplementary Figure 1B,C and D). Since lungs showed the highest level of metastasis in this model, lung tissues were further assessed for metastasis by H&E. We observed that both the number of metastatic foci, as well as the size of the metastatic lesions were significantly reduced in control versus anti-Gas6 treated mice (Figure 1D and E). As a consequence the overall metastatic burden was very significantly reduced in the mice treated with anti-Gas6 blocking antibody compared to control mice (Figure 1F). These data suggest that blockade of Gas6 affects the metastatic cascade at different stages, affecting the metastatic spreading and/or initial seeding as well as the metastatic outgrowth of disseminated pancreatic cancer cells.

**Figure 1.**
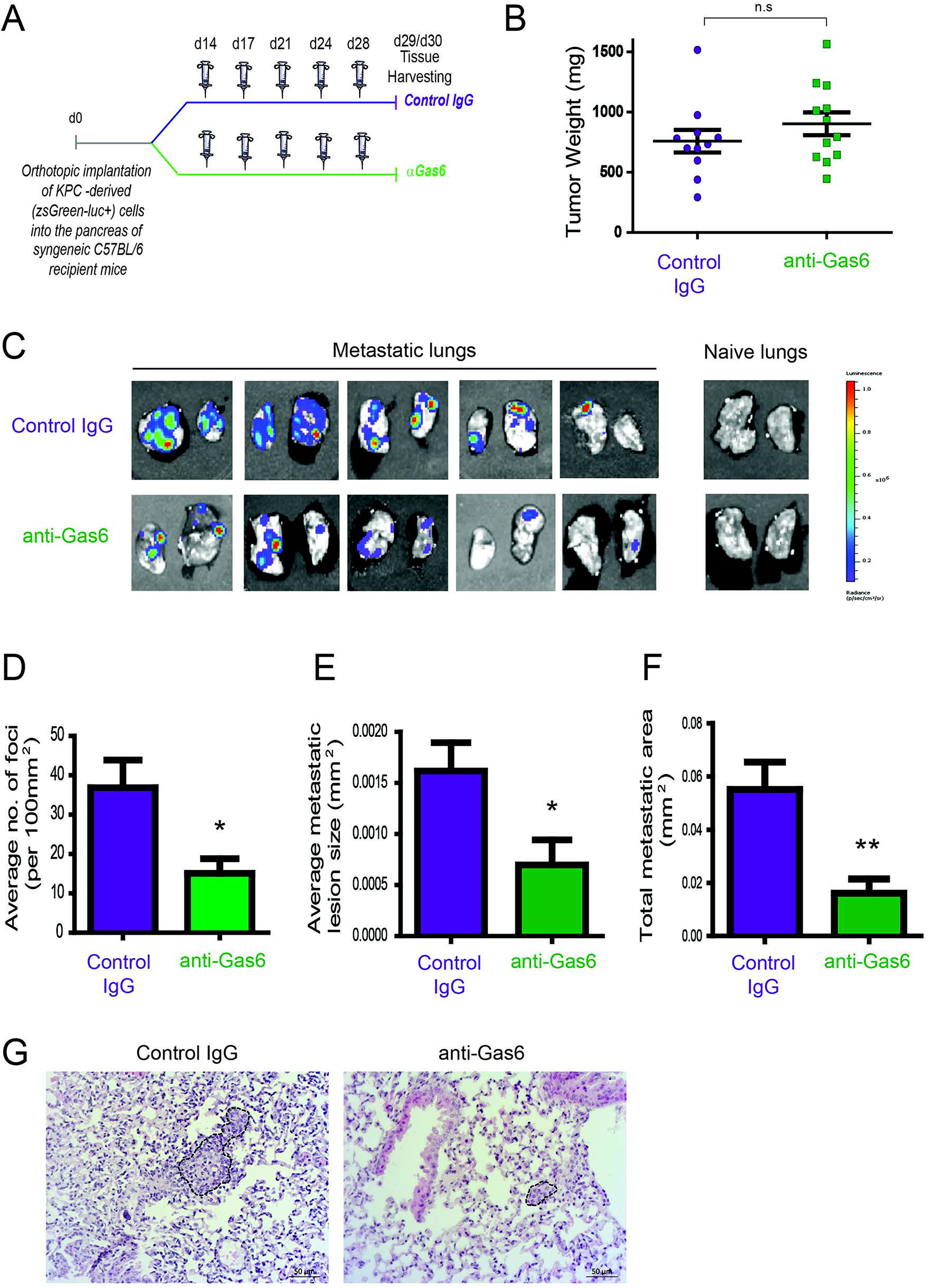
Pharmacological blockade of Gas6 inhibits pancreatic cancer metastasis. **(A)** KPC^luc/zsGreen^ (zsGreen) -derived pancreatic tumor cells (FC1242^luc/zsGreen^) were orthotopically implanted into the pancreas of syngeneic C57BL/6 recipient mice, and mice were treated, starting at day 14 after tumor implantation, twice a week i.p., with either isotype control IgG antibody or Gas6 blocking antibody (0.5 mg/ml?). Primary pancreatic tumors, livers, lungs and mesenteric lymph nodes were harvested at day 30. **(B)** Tumor weights (n= 11 mice for control IgG treatment group; n=12 mice for anti-Gas6 treatment groups). **(C)** Representative IVIS images of metastatic lungs from control IgG and anti-Gas6 treated mice. **(D)** Quantification of number of lung metastatic foci per 100mm^2^ in mice treated with control IgG or anti-Gas6 antibody. * p ≤ 0.05, using unpaired student T test. **(E)** Average size of pulmonary metastatic lesions in mice treated with control IgG or anti-Gas6 antibody. * p ≤ 0.05, using unpaired student T test. **(F)** Quantification of total metastatic burden in mice treated with control IgG or anti-Gas6 antibody. ** p ≤ 0.01, using unpaired student T test. **(G)** Representative images of H&E staining of metastatic lungs from control IgG and anti-Gas6 treated mice. Scale bar 50 μm.

### Tumor associated macrophages and fibroblasts are the main sources of Gas6 in pancreatic cancer

Gas6 is a multifunctional protein that is secreted by different cell types. Gas6 has been shown to be produced by macrophages in pre-malignant lesions of a mammary tumor model [28] and in xenograft and orthotopic models of colon and pancreatic cancer [29]. Gas6 can also be produced by tumor cells [30] and fibroblasts [31]. To determine which cell types produce Gas6 in pancreatic tumors, tumors were harvested at day 23, and tumor cells (CD45−/zsGreen+), non-immune stromal cells (CD45−/zsGreen−), M1-like macrophages (CD45+/F4/80+/CD206−) and M2-like macrophages (CD45+/F4/80+/CD206+) were isolated by flow cytometry (Figure 2A and supplementary Figure 2A, B) and analyzed for the expression *of gas6* (Figure 2B). We found that both F4/80+/CD206+ (M2-like macrophages) and αSMA+ stromal cells (Supplementary Figure 2B) are the main sources of gas6 in pancreatic tumors (Figure 2B). *Ex-vivo*, bone-marrow derived macrophages and pancreatic fibroblasts also produce Gas6 (Figure 2B, C). In agreement with these findings, we observed that tumor areas with activated Axl receptor were often surrounded by TAMs and CAFs (Figure 2C). Analysis of Axl expression and activation in pancreatic cancer patient samples has been correlated with a poor prognosis [21, 22] and Axl activation in cancer cells has been shown to support EMT, cell proliferation, metastasis and drug resistance [5]. While these studies have mainly focused on analyzing the expression and function of Axl on the cancer cells, Axl is also expressed in immune cells, endothelial cells and stromal cells and regulates innate immunity [3, 4], angiogenesis [32–34] and fibrosis [31]. In agreement with this multi-functional role for Axl, we found that Axl is activated in both the tumor and the stromal/immune compartment in biopsies from pancreatic cancer patients (Figure 3A, B).

**Figure 2.**
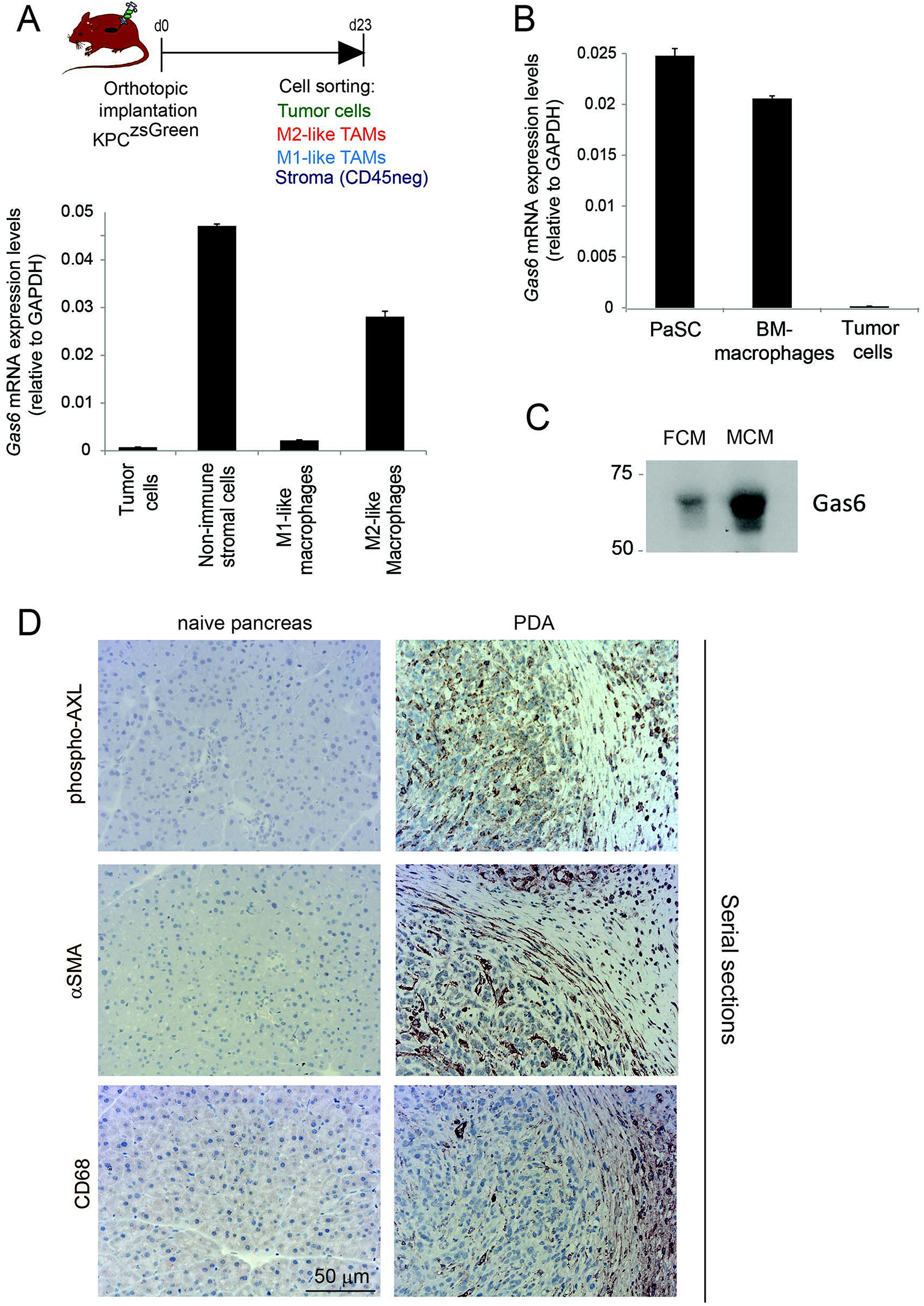
TAMs and CAFs are the main sources of Gas6 in pancreatic tumors. **(A)** KPC^luc/zsGreen^ (zsGreen) -derived tumor cells (FC1242^luc/zsGreen^) were orthotopically implanted into the pancreas of syngeneic recipient (C57/BL6) mice. Tumors were harvested and digested at day 23 after implantation and tumor cells, non-immune stromal cells, M1-like and M2-like macrophages were sorted by flow cytometry. *Gas6* mRNA levels were quantified in CD45-/zsGreen+ tumor cells, CD45-/zsGreen- non-immune stromal cells, CD45+/F4/80+/CD206- M1-like macrophages and CD45+/F4/80+/CD206+ M2-like macrophages isolated from murine pancreatic tumors. Values shown are the mean and SEM (n=3). **(B)** Quantification of *Gas6* mRNA expression levels in mouse primary macrophages and pancreatic fibroblasts. Values shown are the mean and SEM (n=3). **(C)** Immunoblotting analysis of Gas6 secreted protein present in mouse macrophage conditioned media (MCM) and pancreatic fibroblast conditioned media (FCM). **(D)** Images show phospho-Axl, αSMA (fibroblast marker) and CD68 (pan-macrophage marker) staining in naïve mouse pancreas and in serial sections of mouse PDA tissues. Scale bar = 50 μm.

**Figure 3.**
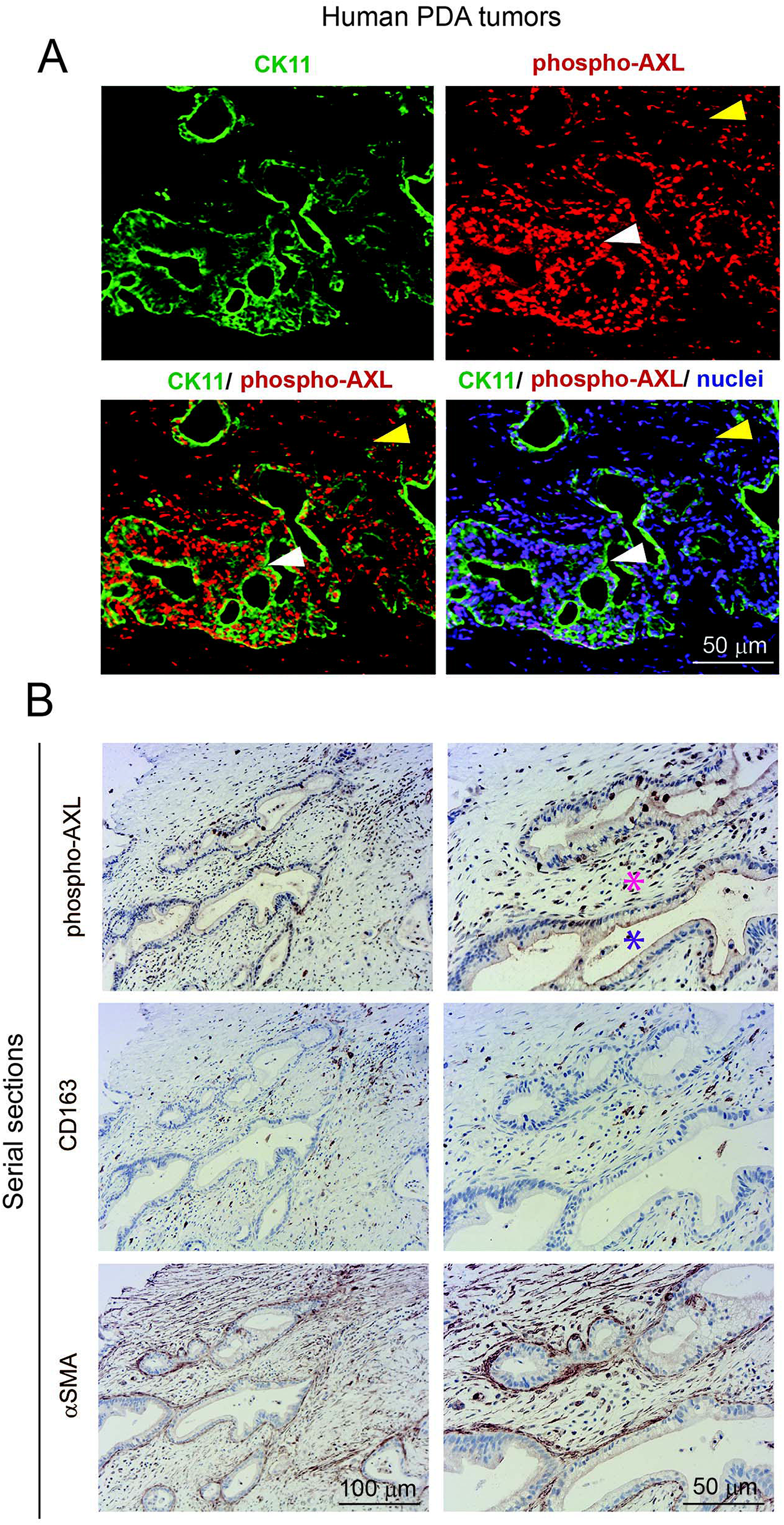
AXL receptor is activated in both the tumor and stromal compartment in biopsies from PDA patients. **(A)** Immunofluorescent staining of human PDA biopsies with CK11 (tumor cell marker, in green), phospho-Axl receptor (in red), and nuclei (in blue). Scale bar, 50 μm. Yellow arrow indicates presence of phosphorylated Axl in the stromal compartment. White arrow indicates presence of phosphorylated Axl in the tumor cells. **(B)** Serial sections of biopsies from human PDA samples immunohistochemically stained for phospho-Axl, CD163 (macrophages) and αSMA (fibroblasts). Scale bars, 50 μm and 100 μm.

### Gas6 blockade alters EMT of pancreatic cancer cells but does not affect angiogenesis or collagen deposition in pancreatic tumors

Previous studies have shown that Gas6-Axl signaling promotes tumor cells’ EMT [35, 36]. To determine whether the reduced metastasis observed when we block Gas6 was caused by an effect on tumor cell EMT we evaluated the expression of EMT markers and transcription factors on tumor cells from pancreatic tumors treated with isotype control antibody or Gas6 blocking antibody. Tumor cells isolated from pancreatic tumors were analysed for the expression of the EMT transcription factors *Snail 1*, *Snail 2*, *Twist 1*, *Twist 2*, *Zeb 1 and Zeb 2* (Figure 4A), the epithelial markers *E-cadherin*, *b-catenin* and *Epcam* and the mesenchymal markers *Vimentin* and *N-cadherin* (Figure 4B). We found that blocking of Gas6 significantly decreased the expression of the EMT transcription factors *Snail 1*, *Snail 2* and *Zeb 2* but did not alter the expression of T*wist 1*, *Zeb 1*, or *Twist 2* (the latter was hardly expressed in these pancreatic cancer cells) (Figure 4A). In agreement with this observation, Gas6 blockade also decreased the expression of the mesenchymal marker *Vimentin*, while *N-cadherin* levels were very low and remained unchanged. However, *E-cadherin* and *B-catenin* levels were also decreased upon anti-Gas6 treatment, suggesting that Gas6 signaling partially regulates cancer cell plasticity [37, 38].

**Figure 4.**
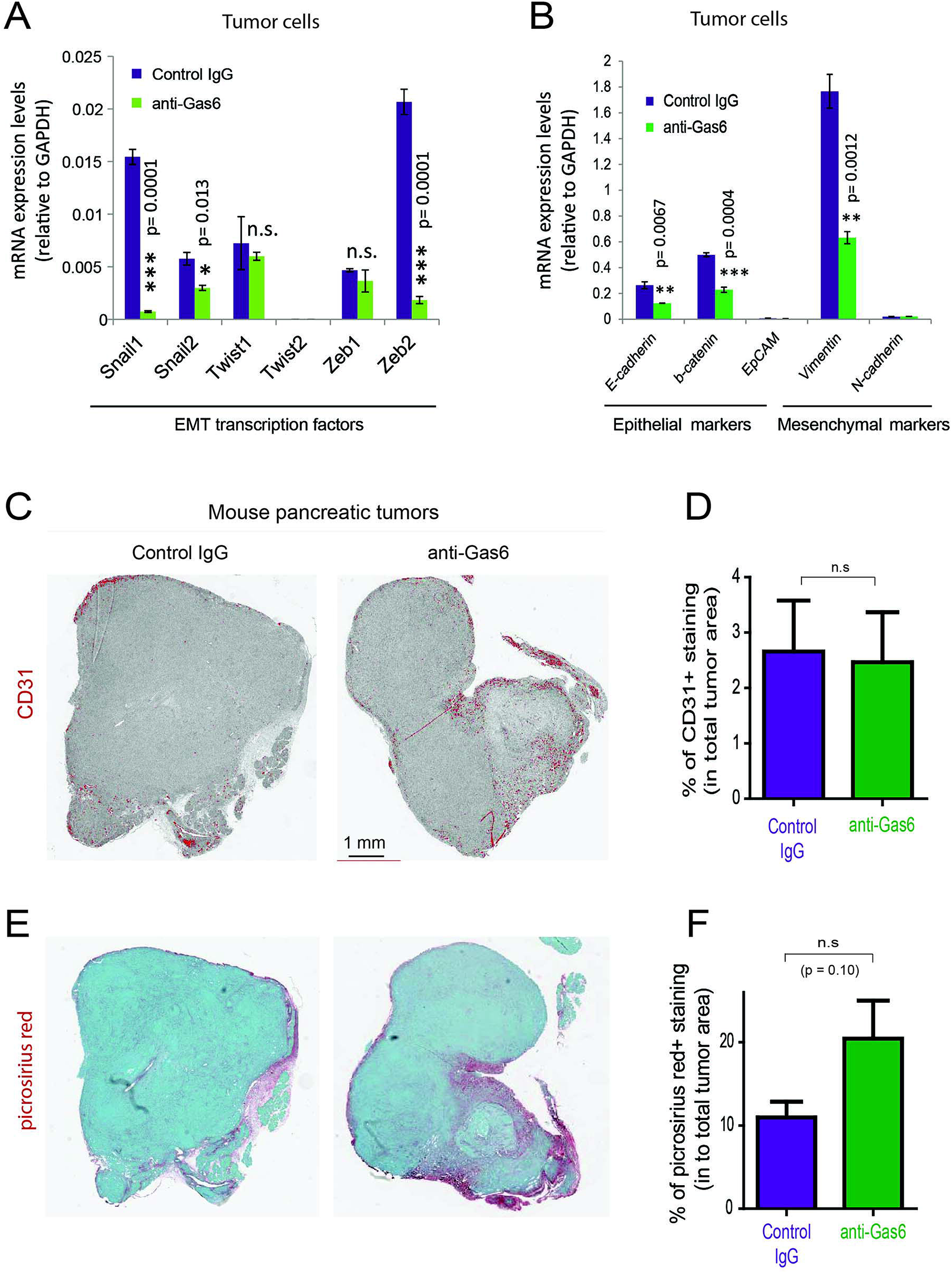
Gas 6 blockade in pancreatic tumors partially affects EMT of tumor cells but does not significantly affect angiogenesis or collagen deposition. **(A)** Quantification of the expression levels of the EMT transcription factors: *Snail 1*, *Snail 2*, *Twist 1*, *Twist 2*, *Zeb 1 and Zeb 2* in tumor cells isolated from mouse PDA tumors. Values shown are the mean and SEM (n=3). **(B)** Quantification of the expression levels of the epithelial markers: *E-cadherin*, *b-catenin*, *EpCAM* and the mesenchymal markers *vimentin and N-cadherin* in tumor cells isolated from mouse PDA tumors. Values shown are the mean and SEM (n=3). * p ≤ 0.05, using unpaired student T test; ** p ≤ 0.01, using unpaired student T test; *** p ≤ 0.005, using unpaired student T test. **(C)** Images of whole scanned pancreatic tumors from control and anti-Gas6 treated mice stained for CD31. **(D)** Quantification of CD31+ staining in total tumor area. Values shown are the mean and SEM (n=3 per treatment group). n.s. no statistically significant differences, using unpaired student T test. **(E)** Images of whole scanned pancreatic tumors from control and anti-Gas6 treated mice stained with picrosirius red. **(F)** Quantification of picrosirius red + staining in total tumor area. Values shown are the mean and SEM (n=4 per treatment group). n.s. no statistically significant differences, using unpaired student T test.

Pancreatic tumors are usually poorly vascularized but since Gas6 signaling can support endothelial cells proliferation and vascularization [33, 39, 40] we next evaluated whether anti-Gas6 therapy could affect angiogenesis in pancreatic tumors. Pancreatic tumor tissues from control and anti-Gas6 treated mice were stained with the endothelial marker CD31, whole tumor tissues were scanned and quantified for CD31 expression which remained unchanged in both treatment groups (Figure 4C, D). Gas6 can also regulate fibroblast proliferation and function. Fourcot et al., showed, in a liver fibrosis model, that Gas6 is secreted by macrophages and fibroblasts and that Gas6 deficiency decreases TGFb and collagen I production by hepatic fibroblasts [31].

Gas6 also stimulates the proliferation of cardiac fibroblasts [41]. Since fibrosis and collagen deposition have been suggested to re-strain the metastatic spreading of pancreatic cancer cells [42–45], we next investigated whether Gas6 blockade could affect collagen deposition in pancreatic tumors. Pancreatic tumor tissues from control and anti-Gas6 treated mice were stained with picrosirius red to assess collagen deposition. Whole tumor tissues were scanned and quantified for collagen deposition (Sirius red positive areas). We observed a slight increase in collagen deposition in tumors from mice treated with anti-Gas6 antibody compared to control but this increase was not statistically significant (Figure 4E, F). These findings suggest that the anti-metastatic effect of Gas6 blockade in pancreatic cancer is not due to changes in angiogenesis or fibrosis.

### Gas6 blockade does not affect myeloid cells or T cells

TAM receptors are also expressed by immune cells and regulate myeloid cell and T-cell functions [3, 46]. Thus, next, with the aim to understand the systemic effect of Gas6 blockade in myeloid cells and T cells in pancreatic cancer, we evaluated the number and activation status of myeloid cells and T cells in pancreatic tumors, blood and metastatic tissues using mass and flow cytometry. Mass cytometry analysis of myeloid (CD11b+) cells, neutrophils/MDSCs (CD11b+/Ly6G+), monocytes (CD11b+/Ly6C+), macrophages (CD11b+/F4/80+), MHC-II+, CD206+ and PD-L1+ macrophages (Figure 5A) and T cells (CD3+), helper T-cells (CD3+/ CD4+), regulatory T cells (CD3+/CD4+/CD25+), cytotoxic T cells (CD3+/CD8+), activated/exhausted cytotoxic T cells (CD8+/CD69+; CD8+/PD-1+) (Figure 5B) from pancreatic tumors from control versus anti-Gas6 treated mice did not show any significant differences (Figure 5A, B and Supplementary Figure 3 A, B). Similarly, myeloid cell and T cell numbers in blood (Supplementary Figure 4A, B) and metastatic lungs from mice treated with control or anti-Gas6 antibody remained the same (Supplementary Figure 5A, B).

**Figure 5.**
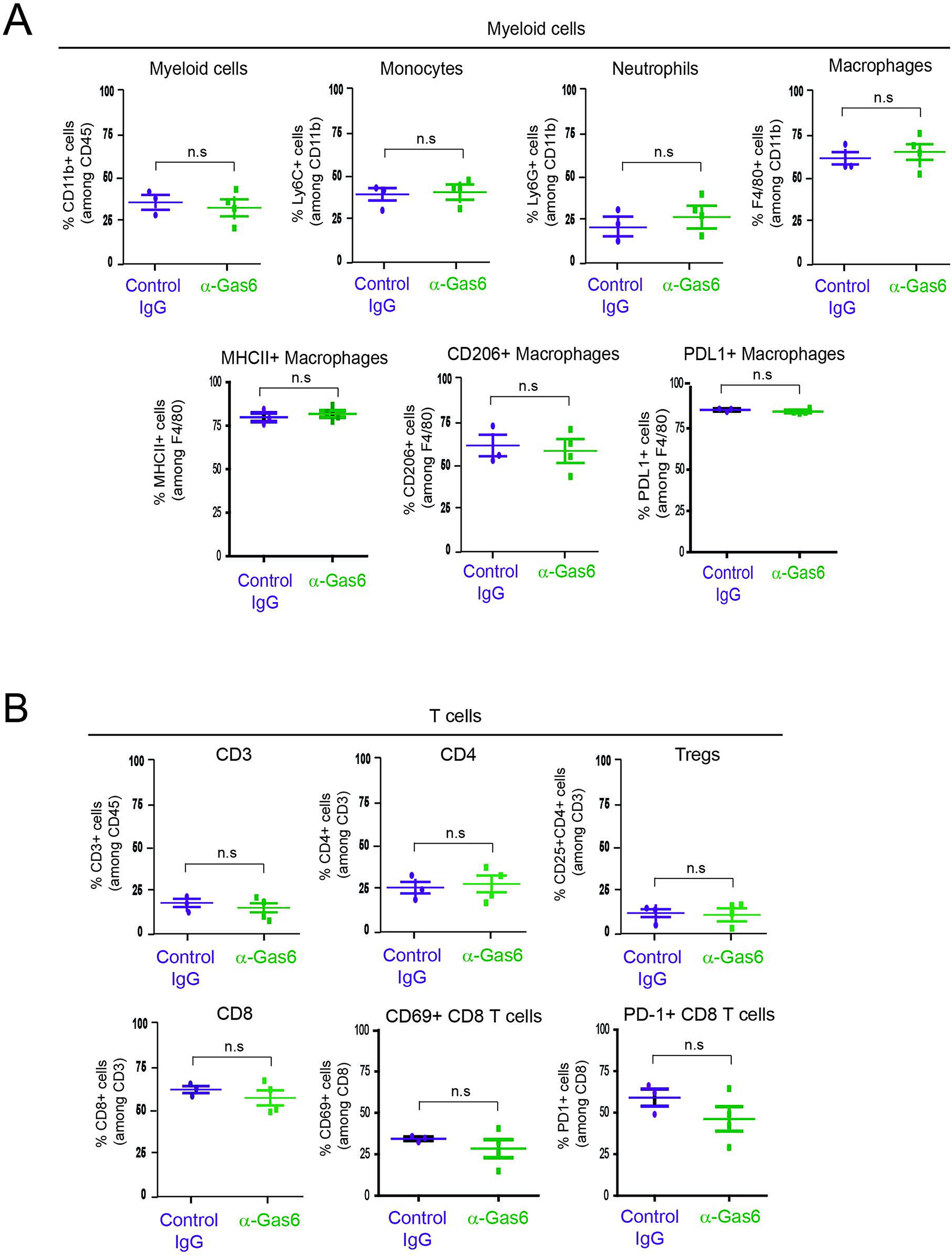
Gas6 blockade does not affect the composition or activation status of myeloid cells and T cells. **(A)** Mass cytometry quantification of CD11b + myeloid cells, Ly6C high/Ly6C low monocytes/MDSCs, Ly6G high/Ly6C low neutrophils/MDSCs, F4/80+ macrophages, MHCII+ macrophages, CD206+ macrophages and PD-L1+ macrophages in mouse pancreatic tumors treated with control IgG (n=3) or anti-Gas6 neutralizing antibody (n=4). **(B)** Mass cytometry quantification of CD3+ T cells, CD4+ T cells, CD4+/CD25+ regulatory T cells (Tregs), CD8+ T cells, CD69+/CD8+ T cells and PD-1+/CD8+ T cells in mouse pancreatic tumors treated with control IgG (n=3) or anti- Gas6 neutralizing antibody (n=4). Graphs were generated with ViSNE data using Cytobank software.

### Gas6 blockade restores NK cell activation and infiltration in metastatic lesions

TAM signaling is involved in the development of natural killer (NK) cells [47]. In an elegant study, Paolino et al., demonstrated that TAM receptor inhibition activates NK cells cytotoxic function and thereby decreases metastasis in mouse models of breast cancer and melanoma [9]. Thus, we next hypothesized that the anti-metastatic effect of Gas6 blockade we observe in our pancreatic cancer model could be due to a re-activation of NK cells. To test this hypothesis we evaluated NK cells in tumor draining lymph nodes, primary pancreatic tumors and metastatic lesions of mice treated with control IgG or anti-Gas6 antibody. The number of NK cells in lung metastatic lesions was significantly higher in mice treated with anti-Gas6 antibody compared to control treated mice (Figure 6 A, B). The number of NK cells, and in particular the number of proliferating NK cells, was also increased in tumor draining lymph nodes from anti-Gas6 treated mice compared to control treated mice (Figure 6 C, D). However, NK cells were almost absent in all primary tumors from both anti-Gas6 and control treated mice (except for one anti-Gas6 treated pancreatic tumor). (Supplementary Figure 6).

**Figure 6.**
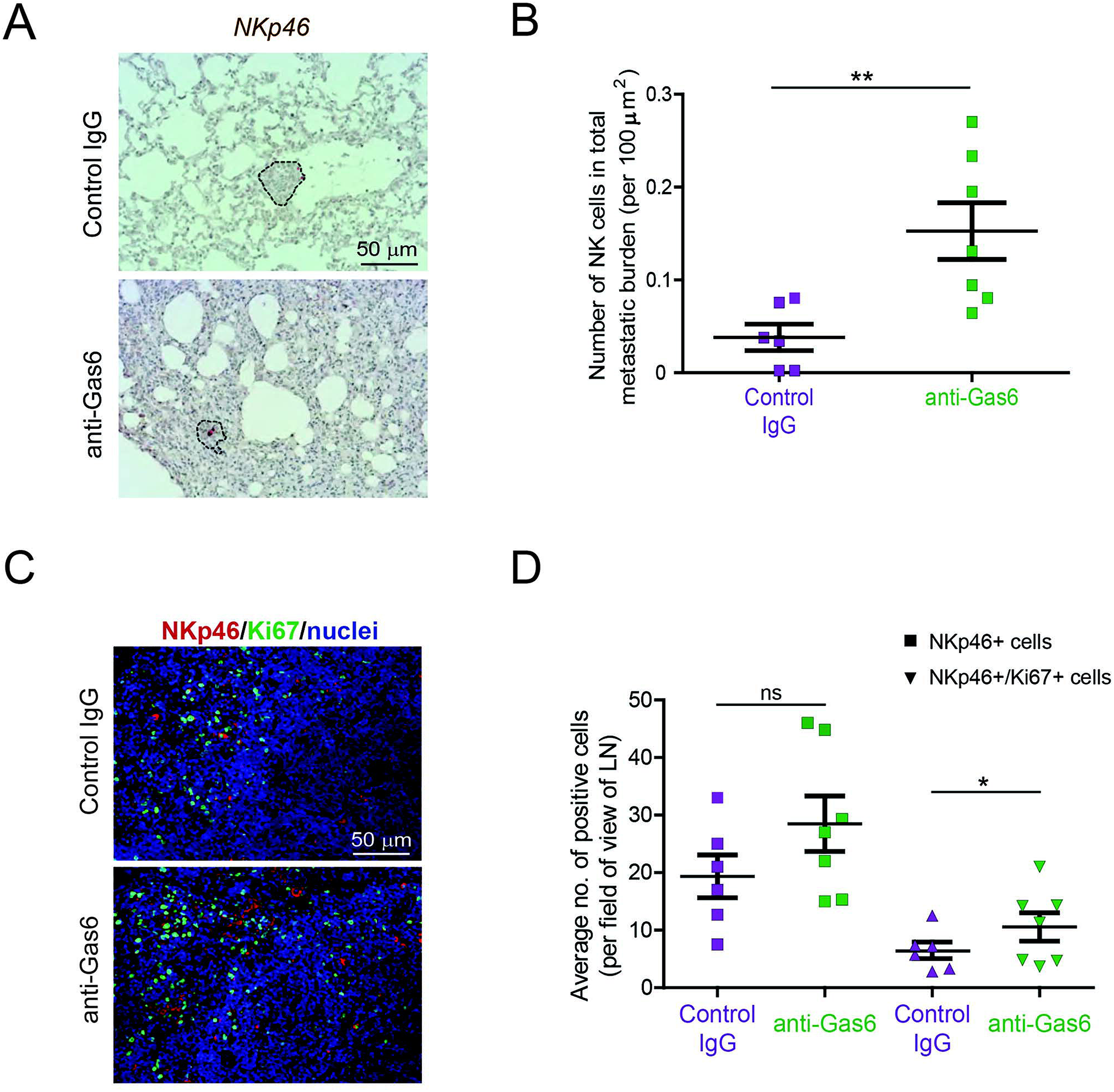
Gas6 blockade increases NK cell numbers in metastatic lungs and in tumor draining lymph nodes. **(A)** Immunohistochemical staining of NK cells in metastatic lungs from pancreatic tumor bearing mice treated with control IgG or anti-Gas6 antibody. Lesions indicated by dashed line and NK cells by red asterisk. Scale bar, 50 μm. **(B)** Quantification of NK cells in metastatic lung tissues from control IgG and anti-Gas6 treated mice. Values shown are the mean and SEM (n=6 mice in IgG treatment group, n=7 mice in anti-Gas6 treatment group). ** p ≤ 0.01, using unpaired student T test. **(C)** Immunofluorescent staining of NK cells in mesenteric lymph nodes from pancreatic tumor bearing mice treated with control IgG or anti-Gas6 antibody. NK marker NKp46 is shown in red, Ki67 is shown in green and nuclei were stained with DAPI (in blue). Scale bar, 50 μm. **(D)** Quantification of NK cells in tumor draining lymph nodes from control IgG and anti-Gas6 treated mice. Values shown are the mean and SEM (n=6 mice IgG treatment group and n=7 mice anti-Gas6 treatment group, 3-6 fields/ mouse tissue were quantified). * p ≤ 0.05, using unpaired student T test.

## Discussion

The data presented in this study describe a dual anti-tumor effect of Gas6 blockade in pancreatic tumors, shedding light on the anti-cancer mechanism of action of inhibitors of the Gas6-Axl pathway and supporting the rationale for using anti-Gas6 therapy in pancreatic cancer patients. In these studies we show that blockade of Gas6 in pancreatic tumors acts simultaenously on both the tumor cells, reversing their plasticity, as well as on NK cells, promoting their activation and recruitment to the metastatic site (Figure 7). This dual function of anti-Gas6 therapy explains our observation that Gas6 blockade inhibits both the number and the size of the metastatic lesions. These findings suggest that anti-Gas6 therapy impairs several steps of the metastatic cascade including the initial spreading of tumor cells and the metastatic outgrowth of disseminated pancreatic cancer cells by acting on both the tumor cells and the NK cells.

**Figure 7.**
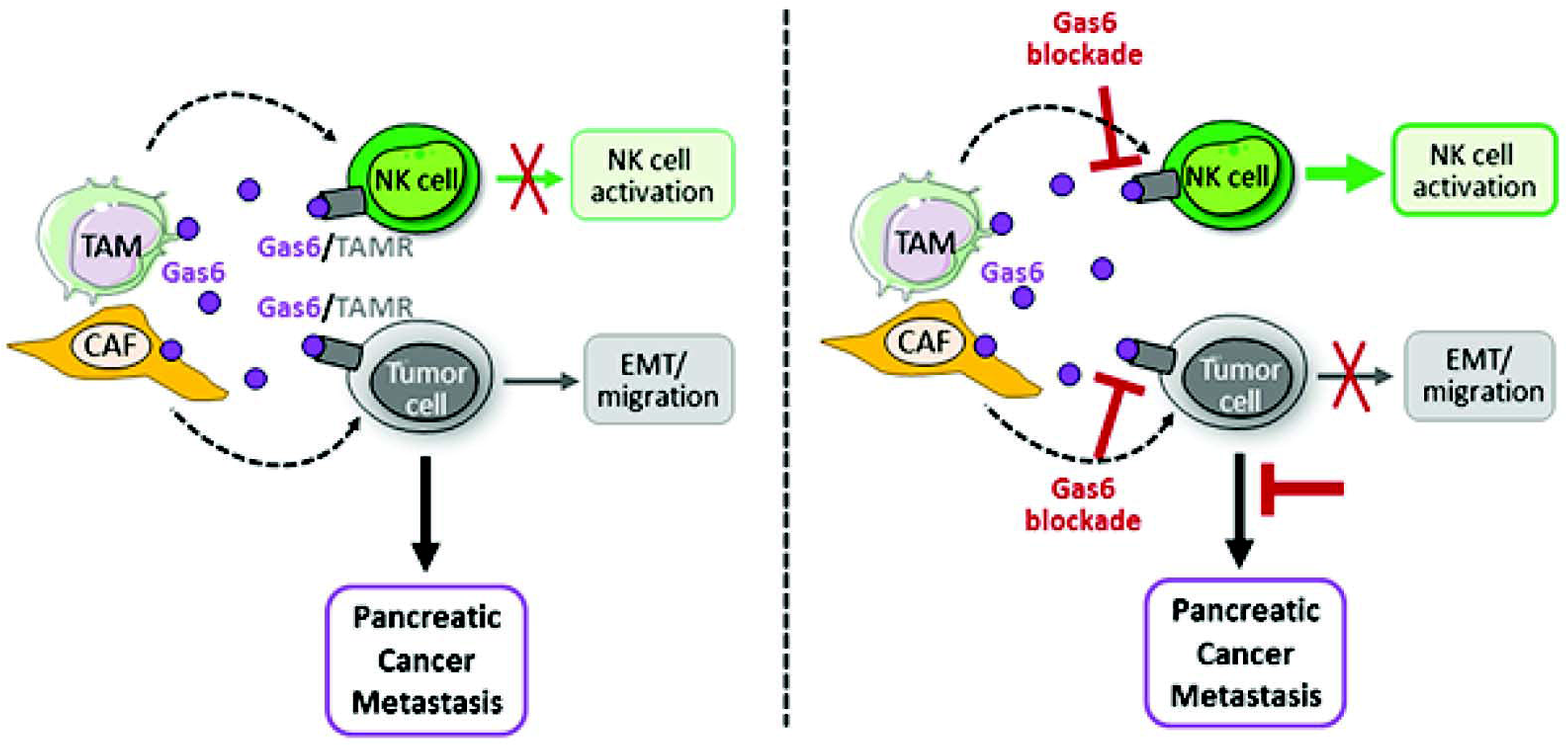
Schematics depicting the multifunctional role of stroma-derived Gas6 in pancreatic cancer. Gas6 produced by tumor associated macrophages (TAMs) and cancer associated fibroblasts (CAFs) in pancreatic tumors binds to Tyro-Axl-Mer (TAM) receptors expressed on the surface of both tumor cells and NK cells, and thereby acts on both: the tumor cells by EMT and metastasis and the NK cells by inactivating their anti-tumor function. *In vivo* blockade of Gas6 partially reverses tumor cells EMT and re-activates NK cells, leading to a decrease in pancreatic cancer metastasis.

So far many studies have focused on the cancer-cell autonomous role of Gas6 and based on their effect on tumor cell proliferation and plasticity several inhibitors of the Gas6-Axl pathways are currently being tested in pancreatic cancer patients.

Our studies show that Gas6 inhibition in pancreatic cancer not only affects the tumor cells but also the NK cells, thus suggesting that the activation status of NK cells should also be assessed in patients and could be used as a biomarker to monitor response to Gas6/Axl inhibitors.

Gas6/Axl signaling is a negative regulator of the immune system and inhibition of the Gas6-Axl signaling leads to autoimmunity [4]. While the function of Gas6-Axl signaling on tumor cell proliferation, EMT, migration and drug resistance has been extensively studied [5], only a few studies have investigated the role of Gas6/Axl signaling in the immune system in the context of cancer [7, 9, 24]. Guo et al., found that the Axl inhibitor R428 inhibited tumor growth of subcutaneously implanted murine 4T1 breast cancer cells and intra-peritoneally implanted murine ID8 ovarian cancer cells by activating CD4+ and CD8+ T cells [7]. Inspired by this study, we investigated whether, in our pancreatic cancer model, Gas6 blockade supports the activation of T cells. Unlike Guo et al., we did not observe any statistically significant difference in CD4+ or CD8+ T cells in pancreatic tumors, blood or metastatic tissues, in control versus anti-Gas6 treated mice. Ludwig et al., found that treating mouse pancreatic tumors with the Axl inhibitor BGB324 decreased the number of tumor associated macrophages (TAMs) [24]. However, in our study, blocking Gas6 did not affect TAMs or other myeloid cell populations in primary tumors, blood or metastatic organs. Together, these results suggest that Axl inhibition may act on the immune system differently to Gas6 blockade affecting T cells or macrophages versus NK cells. In another study, Paolino et al., showed that TAM receptor inhibition activates NK cells in mouse tumor models of melanoma and breast cancer leading to decreased tumor growth [9]. In agreement with these findings, we found that blocking Gas6 in mice bearing pancreatic tumors, increases NK cell activation in tumor draining lymph nodes and NK cell recruitment to the metastatic site, and decreases pancreatic cancer metastasis.

Inhibition of the Gas6-Axl pathway has been shown to reverse EMT, tumor migration and intra-tumoral microvessel density in pancreatic cancer [23]. In agreement with these findings, we found that Gas6 blockade indeed partially reverses EMT of pancreatic cancer cells, however, we did not observe any significant changes in tumor angiogenesis in our model. Pancreatic tumors are usually hypo-vascularized compared to a normal pancreas and anti-angiogenic therapies have not been successful in pancreatic cancer [48]. Similar to the human disease, in our pancreatic mouse tumor model, tumors are poorly vascularized and blocking Gas6 did not show any further decrease in tumor vascularization. Loges et al., previously showed that tumor associated macrophages (TAMs) produce Gas6 in various mouse tumor models [11]. In our study we find that both TAMs and CAFs are the main sources of Gas6 in pancreatic tumors. These findings suggest that the abundance of TAMs and CAFs in pancreatic cancer patients could be used to determine which patients would benefit the most from anti-Gas6 therapy.

In conclusion, our studies suggest that in pancreatic cancer, Gas6 is secreted by both TAMs and CAFs and blockade of Gas6 has a dual anti-metastatic effect by acting on both the tumor cells and the NK cells. Thus, inactivation of Gas6 signaling can promote anti-tumor immunity, via NK cell activation, in pancreatic tumors. Since this Gas6-dependent immune regulation of NK cells is also conserved in humans, anti-Gas6-Axl therapies are likely to promote anti-tumor immunity, via NK cell activation, in pancreatic cancer patients. This study provides further mechanistic insights into the mode of action of anti-Gas6 therapy and suggests the use of NK cells as an additional biomarker for response to anti-Gas6 therapy in pancreatic cancer patients.

## Materials and Methods

### Generation of primary KPC-derived pancreatic cancer cells

The murine pancreatic cancer cells KPC FC1242 were generated in the Tuveson lab (Cold Spring Harbor Laboratory, New York, USA) isolated from pancreatic ductal adenocarcinoma (PDA) tumor tissues obtained from LSL-Kras^G12D^; LSL-Trp53^R172H^; Pdx1-Cre mice of a pure C57BL/6 background as described previously with minor modifications [49].

### Generation of primary macrophages, primary pancreatic fibroblasts, macrophage (MCM) and fibroblasts (FCM) conditioned media

Primary murine macrophages were generated by flushing the bone marrow from the femur and tibia of 6-8 week old C57BL/6 mice followed by incubation for 5 days in DMEM containing 10% FBS and 10 ng/mL murine M-CSF (Peprotech). Primary pancreatic stellate cells were isolated from the pancreas of C57BL/6 mice by density gradient centrifugation, and were cultured on uncoated plastic dishes in IMDM with 10% FBS and 4mM L-glutamine. Under these culture conditions pancreatic stellate cells activated into myofibroblasts.

To generate macrophage and fibroblast conditioned media, cells were cultured in serum free media for 24-36 h, supernatant was harvested, filtered with 0.45◻m filter, concentrated using StrataClean Resin (Agilent Technologies) and immunoblotted for Gas6 (R&D Systems, AF885).

### RTK arrays & immunoblotting

Cells were serum starved or treated with macrophage conditioned media for 30 min or 3h, harvested and lysed in RIPA buffer (150 mM NaCl, 10 mM Tris-HCl pH 7.2, 0.1% SDS, 1% Triton X-100, 5 mM EDTA) supplemented with a complete protease inhibitor mixture (SIGMA), a phosphatase inhibitor cocktail (Invitrogen), 1 mM PMSF and 0.2 mM Na_3_VO_4_. Cell lysates were analyzed with the Phospho-RTK Array Kit (R&D Systems). Immunoblotting analyses was performed using phospho-Axl antibody (R&D systems, AF2228), Axl antibody (R&D systems, AF854 ref) and tubulin antibody (Sigma, T6199) as loading control.

### Syngeneic Orthotopic pancreatic cancer model

1 × 10^6^ primary KPC^luc/zsGreen^ cells (FC1242^luc/zsGreen^) isolated from a pure C57Bl/6 background were implanted into the pancreas of immune-competent syngeneic C57Bl/6 six- to eight-week-old female mice, and tumors were established for two weeks before beginning treatment. Mice were administered i.p with Gas6 neutralizing antibody (2 mg/kg), or IgG isotype control antibody, every 3 −4 days for 15 days before harvest.

### Analysis and quantification of immune cells in pancreatic tumors by mass cytometry

Pancreatic tumors were resected from the mice and mechanically and enzymatically digested in Hanks Balanced Salt Solution (HBSS) with 1 mg/mL Collagenase P (Roche) Cell suspensions were centrifuged for 5 min at 1500 rpm, resuspended in HBSS and filtered through a 500 μm polypropylene mesh (Spectrum Laboratories). Cells were resuspended in 1 mL 0.05%Trypsin and incubated at 37°C for 5 minutes. Cells were filtered through a 70 μm cell strainer and resuspended in Maxpar cell staining buffer (Fluidigm). The samples were centrifuged for 5 min at 450 *x g* and supernatant removed. The cells were subsequently stained with Cell-ID 195-Cisplatin (Fluidigm) viability marker diluted 1:40 in Maxpar PBS (Fluidigm) for 5 min. Cells were centrifuged at 450 *x g* for 5 min and washed twice in Maxpar cell staining buffer. Samples were blocked for 10 minutes on ice with 1:100 diluted FC Block (BD Pharmingen, Clone 2.4G2) and metal-conjugated antibody cocktail added and incubated for 30 min at 4°C. Antibodies were used at the concentrations recommended by manufacturers. Cells were washed twice in cell staining buffer and stained with 125 μM 191-Intercalator-Ir (Fluidigm) diluted in 1:2000 Maxpar fix and perm buffer (Fluidigm) overnight at 4 °C. The cells were washed twice in Maxpar cell staining buffer and centrifuged at 800 *x g* for 5 min. A post-fix was performed by incubating the cells in 1.6% PFA for 30 min at RT. Cells were washed twice in 18Ω distilled water (Fluidigm), mixed 1:10 with EQTM Four Element Calibration Beads (Fluidigm) and acquired on the Helios CyTOF system (Fluidigm). Samples were acquired at a rate of around 200 cells/s. All generated FCS files were normalized and beads removed [50]. All analysis was performed in Cytobank: Manual gating was used to remove dead cells (195Pt+) and debris and to identify single cells (191 Ir+).

viSNE analysis was performed on the data utilising t-stochastic neighbour embedding (t-SNE) mapping based on high dimensional relationships. CD45+ population selected by manual gating was used as the starting cell population and using proportional sampling viSNE unsupervised clustering was performed. Manual gating was then performed on the viSNE map created to determine cell population percentages. Spanning-tree Progression Analysis of Density-normalized Events (SPADE) analysis was performed in Cytobank using manual gated CD45+ cells, 200 target number of nodes and 10% down sampled events, to equalize the density in different parts of the cloud. Gating of cell populations was performed to identify major cell populations and percentages.

### FACS sorting and analysis of blood and lungs by flow cytometry

Single cell suspensions from murine primary pancreatic tumors and pulmonary metastasis were prepared by mechanical and enzymatic disruption and tumor cells, tumor associated macrophages and stromal cells were analysed and sorted using flow cytometry (FACS ARIA II, BD Bioscience). Samples were digested as outlined above, the cells were then filtered through a 70 μm cell strainer and resuspended in PBS + 1% BSA, blocked for 10 minutes on ice with FC Block (BD Pharmingen, Clone 2.4G2) and stained with Sytox® blue viability marker (Life Technologies) and conjugated antibodies anti-CD45-PE/Cy7 (Biolegend, clone 30-F11) and anti-F4/80-APC (Biolegend, clone BM8).

Blood was collected from mice via tail vein bleed in EDTA-tubes. Red blood cell lysis was performed and resulting leukocytes were resuspended in PBS + 1% BSA and blocked for 10 mins on ice with FC block and stained with Sytox® blue viability marker and conjugated antibodies anti-CD45-APC/Cy7 (Biolegend, 103115), anti-CD11b-APC (Biolegend, 101212), anti-Ly6G-PerCP-Cy5.5 (Biolegend, 127616), anti-Ly6C-PE (Biolegend, 128008), anti-CD3-PE-Cy7 (Biolegend, 100320), anti-CD4-PE (Biolegend, 100408) and anti-CD8-PerCP-Cy5.5 (Biolegend, 100734). Cell analysis was performed using FACS Canto II.

### Gene expression

Total RNA was isolated from FACS sorted tumor cells, tumor associated macrophages and non-immune stromal cells from primary pancreatic tumors as described in Qiagen Rneasy protocol. Total RNA from the different cell populations was extracted using a high salt lysis buffer (Guanidine thiocynate 5 M, sodium citrate 2.5 uM, lauryl sarcosine 0.5% in H2O) to improve RNA quality followed by purification using Qiagen Rneasy protocol. cDNA was prepared from 1μg RNA/sample, and qPCR was performed using gene specific QuantiTect Primer Assay primers from Qiagen. Relative expression levels were normalized to *gapdh* expression according to the formula <2^− (Ct *gene of interest* − Ct *gapdh*) [51]

### Quantification of metastasis

#### By IVIS imaging

IVIS spectral imaging of bioluminescence was used for orthotopically implanted tumor cells expressing firefly luciferase using IVIS spectrum system (Caliper Life Sciences). Organs were resected for *ex vivo* imaging coated in 100 μL D-luciferin (Perkin Elmer) for 1 min and imaged for 2 min at automated optimal exposure. Analysis was performed on the Living Image software (PerkinElmer) to calculate the relative bioluminescence signal from photon per second mode normalised to imaging area (total flux) as recommended by the manufacturer.

#### By H&E staining

FFPE lungs were serially sectioned through the entire lung using microtome at 4 μm thickness. Sections were stained with H&E and images were taken using a Zeiss Observer Z1 Microscope (Zeiss) to identify metastatic foci. The number of foci were counted, and the total area of metastatic foci was measured using Zen imaging software. Metastatic burden was calculated by the following equations:

No. of foci per 100 mm^2^: *(Average no. foci per section/ average tissue area per section (mm^2^))* \**100*
Average metastatic lesion size (mm^2^): *Average total area of metastasis (mm^2^)/ average number of foci per section*
Total metastatic burden: *Sum of area of each foci of each section*

### Immunohistochemistry and Immunofluorescence

Deparaffinization and antigen retrieval was performed using an automated DAKO PT-link. Paraffin-embedded pancreatic tumors, lymph nodes and lung metastasis tissues were immuno-stained using the DAKO envision+ system-HRP.

#### Antibodies and procedure used for Immunohistochemistry

All primary antibodies were incubated for 2 hours at room temperature: αSMA (Abcam, ab5694 used at 1:200 after low pH antigen retrieval), CD31 (Cell signalling technology, CST 77699 used at 1:100 after low pH antigen retrieval), NKp46 (Biorbyt, orb13333 used at 1:200 after low pH antigen retrieval), CD3 (Abcam, ab5690 used at 1:100 after high pH antigen retrieval), CD68 (Abcam, ab31630 used at 1:400 after low pH antigen retrieval) and CD206 (Abcam, ab8918 used at 1:400 after low pH antigen retrieval). Subsequently, samples were incubated with secondary HRP-conjugated antibody (from DAKO envision kit) for 30 min at room temperature. All antibodies were prepared in antibody diluent from Dako envision kit. Staining was developed using diamino-benzidine and counterstained with hematoxylin.

Human paraffin-embedded PDA tissue sections were incubated overnight at RT with the following primary antibodies: phospho-Axl (R&D, AF2228, used 1:500 after high pH antigen retrieval), CD163 (Abcam, ab74604 pre-diluted after low pH antigen retrieval), αSMA (Abcam, ab5694 used 1:100 after low pH antigen retrieval),

#### Antibodies and procedure used for Immunofluorescence

After low pH antigen retrieval, lymph node tissue sections derived from mice bearing pancreatic tumors were incubated overnight at RT with the following primary antibodies: NKp46 (R&D systems AF2225, used at 1:25), Ki67 (Abcam ab15580, used at 1:1000). Samples were washed with PBS and incubated with donkey anti-goat 594 (Abcam ab150132) and donkey anti-rabbit 488 (Abcam ab98473) secondary antibodies respectively, all used at 1:300 and DAPI at 1:600 for 2 hours at RT. Slides were washed with PBS, final quick wash with distilled water and mounted using DAKO fluorescent mounting media.

Human PDA frozen tissue sections were fixed with cold acetone, permeabilized in 0.1% Triton, blocked in 8% goat serum and incubated overnight at 4°C with anti-phospho Axl (R&D, AF2228, diluted 1:200) CK11 (Cell signaling, CST 4545, diluted 1:200), followed by fluorescently labelled secondary antibodies goat anti mouse 488 (Abcam ab98637), goat anti-rabbit 594 (Abcam ab98473) used at 1:300 for 2 hours at RT slides were washed with PBS, final quick wash with distilled water and mounted using DAKO fluorescent mounting media.

### Statistical Methods

Statistical significance for *in vitro* assays and animal studies was assessed using unpaired two-tailed Student *t* test and the GraphPad Prism 5 program. All error bars indicate SD of n=3 (*in vitro* studies) or SEM n= 7-8 (animal studies).

### Institutional approvals

All studies involving human tissues were approved by the University of Liverpool and were considered exempt according to national guidelines. Human pancreatic cancer samples were obtained from the Liverpool Tissue Bank from patients that consented to use the surplus material for research purposes. All animal experiments were performed in accordance with current UK legislation under an approved project licence (reference number: 403725). Mice were housed under specific pathogen-free conditions at the Biomedical Science Unit at the University of Liverpool.

## AUTHOR CONTRIBUTIONS

L.I designed experiments and performed most of the experiments including *in vivo* experiments, mass cytometry/flow cytometry, cell isolations, immunohistochemical stainings and qPCR experiments. T. L designed experiments, helped with tissue harvesting, tissue stainings, primary cell isolations and qPCR experiments. A.M. designed experiments, helped with tissue harvesting and tissue stainings. M.C.S. provided conceptual advice and help with *in vivo* experiments. A.M and L.I wrote the manuscript. A.M. conceived and supervised the project. All authors helped with the analysis and interpretation of the data, the preparation of the manuscript, and approved the manuscript.

## Acknowledgments

We thank David Tuveson and Danielle Engle for providing the mouse KPC-derived pancreatic cancer cells. We thank Arthur Taylor and Patricia Murray for transducing the KPC cells with zsGreen/luciferase lentivirus. We thank Almudena Santos for technical support with the immunohistochemistry. We also acknowledge the Liverpool Tissue Bank for providing tissue samples, the mass/flow cytometry/cell sorting facility, the biomedical science unit and the pre-clinical *in vivo* imaging facility for provision of equipment and technical assistance. We thank the patients and their families who contributed with tissue samples to these studies.

## References

1 Paolino M, Penninger JM. The Role of TAM Family Receptors in Immune Cell Function: Implications for Cancer Therapy. Cancers (Basel) 2016; 8.

2 Sasaki T, Knyazev PG, Clout NJ, Cheburkin Y, Gohring W, Ullrich A et al. Structural basis for Gas6-Axl signalling. EMBO J 2006; 25: 80–87.

3 Lemke G, Rothlin CV. Immunobiology of the TAM receptors. Nature reviews Immunology 2008; 8: 327–336.

4 Lu Q, Lemke G. Homeostatic regulation of the immune system by receptor tyrosine kinases of the Tyro 3 family. Science 2001; 293: 306–311.

5 Wu G, Ma Z, Hu W, Wang D, Gong B, Fan C et al. Molecular insights of Gas6/TAM in cancer development and therapy. Cell Death Dis 2017; 8: e2700.

6 Hafizi S, Dahlback B. Gas6 and protein S. Vitamin K-dependent ligands for the Axl receptor tyrosine kinase subfamily. FEBS J 2006; 273: 5231–5244.

7 Guo Z, Li Y, Zhang D, Ma J. Axl inhibition induces the antitumor immune response which can be further potentiated by PD-1 blockade in the mouse cancer models. Oncotarget 2017; 8: 89761–89774.

8 Myers KV, Amend SR, Pienta KJ. Targeting Tyro3, Axl and MerTK (TAM receptors): implications for macrophages in the tumor microenvironment. Molecular cancer 2019; 18: 94.

9 Paolino M, Choidas A, Wallner S, Pranjic B, Uribesalgo I, Loeser S et al. The E3 ligase Cbl-b and TAM receptors regulate cancer metastasis via natural killer cells. Nature 2014; 507: 508–512.

10 Rankin EB, Fuh KC, Castellini L, Viswanathan K, Finger EC, Diep AN et al. Direct regulation of GAS6/AXL signaling by HIF promotes renal metastasis through SRC and MET. Proc Natl Acad Sci U S A 2014; 111: 13373–13378.

11 Loges S, Schmidt T, Tjwa M, van Geyte K, Lievens D, Lutgens E et al. Malignant cells fuel tumor growth by educating infiltrating leukocytes to produce the mitogen Gas6. Blood 2010; 115: 2264–2273.

12 Ireland L, Santos A, Campbell F, Figueiredo C, Hammond D, Ellies LG et al. Blockade of insulin-like growth factors increases efficacy of paclitaxel in metastatic breast cancer. Oncogene 2018.

13 Nielsen SR, Quaranta V, Linford A, Emeagi P, Rainer C, Santos A et al. Macrophage-secreted granulin supports pancreatic cancer metastasis by inducing liver fibrosis. Nature cell biology 2016; 18: 549–560.

14 Qian BZ, Zhang H, Li J, He T, Yeo EJ, Soong DY et al. FLT1 signaling in metastasis-associated macrophages activates an inflammatory signature that promotes breast cancer metastasis. The Journal of experimental medicine 2015; 212: 1433–1448.

15 DeNardo DG, Brennan DJ, Rexhepaj E, Ruffell B, Shiao SL, Madden SF et al. Leukocyte complexity predicts breast cancer survival and functionally regulates response to chemotherapy. Cancer Discov 2011; 1: 54–67.

16 Ireland L, Santos A, Ahmed MS, Rainer C, Nielsen SR, Quaranta V et al. Chemoresistance in Pancreatic Cancer Is Driven by Stroma-Derived Insulin-Like Growth Factors. Cancer Res 2016; 76: 6851–6863.

17 Shree T, Olson OC, Elie BT, Kester JC, Garfall AL, Simpson K et al. Macrophages and cathepsin proteases blunt chemotherapeutic response in breast cancer. Genes Dev 2011; 25: 2465–2479.

18 Siegel RL, Miller KD, Jemal A. Cancer statistics, 2019. CA: a cancer journal for clinicians 2019; 69: 7–34.

19 Ireland LV, Mielgo A. Macrophages and Fibroblasts, Key Players in Cancer Chemoresistance. Front Cell Dev Biol 2018; 6: 131.

20 Zambirinis CP, Miller G. Cancer Manipulation of Host Physiology: Lessons from Pancreatic Cancer. Trends Mol Med 2017; 23: 465–481.

21 Song X, Wang H, Logsdon CD, Rashid A, Fleming JB, Abbruzzese JL et al. Overexpression of receptor tyrosine kinase Axl promotes tumor cell invasion and survival in pancreatic ductal adenocarcinoma. Cancer 2011; 117: 734–743.

22 Koorstra JB, Karikari CA, Feldmann G, Bisht S, Rojas PL, Offerhaus GJ et al. The Axl receptor tyrosine kinase confers an adverse prognostic influence in pancreatic cancer and represents a new therapeutic target. Cancer Biol Ther 2009; 8: 618–626.

23 Kirane A, Ludwig KF, Sorrelle N, Haaland G, Sandal T, Ranaweera R et al. Warfarin Blocks Gas6-Mediated Axl Activation Required for Pancreatic Cancer Epithelial Plasticity and Metastasis. Cancer Res 2015; 75: 3699–3705.

24 Ludwig KF, Du W, Sorrelle NB, Wnuk-Lipinska K, Topalovski M, Toombs JE et al. Small-Molecule Inhibition of Axl Targets Tumor Immune Suppression and Enhances Chemotherapy in Pancreatic Cancer. Cancer Res 2018; 78: 246–255.

25 Gay CM, Balaji K, Byers LA. Giving AXL the axe: targeting AXL in human malignancy. Br J Cancer 2017; 116: 415–423.

26 Pylayeva-Gupta Y, Das S, Handler JS, Hajdu CH, Coffre M, Koralov SB et al. IL35-Producing B Cells Promote the Development of Pancreatic Neoplasia. Cancer Discov 2016; 6: 247–255.

27 Zhu Y, Knolhoff BL, Meyer MA, Nywening TM, West BL, Luo J et al. CSF1/CSF1R blockade reprograms tumor-infiltrating macrophages and improves response to T-cell checkpoint immunotherapy in pancreatic cancer models. Cancer Res 2014; 74: 5057–5069

28 Gomes AM, Carron EC, Mills KL, Dow AM, Gray Z, Fecca CR et al. Stromal Gas6 promotes the progression of premalignant mammary cells. Oncogene 2019; 38: 2437–2450.

29 Loges S, Schmidt T, Tjwa M, van Geyte K, Lievens D, Lutgens E et al. Malignant cells fuel tumor growth by educating infiltrating leukocytes to produce the mitogen Gas6. Blood 2010; 115: 2264–2273.

30 Baumann C, Ullrich A, Torka R. GAS6-expressing and self-sustaining cancer cells in 3D spheroids activate the PDK-RSK-mTOR pathway for survival and drug resistance. Molecular Oncology 2017; 11: 1430–1447.

31 Fourcot A, Couchie D, Chobert MN, Zafrani ES, Mavier P, Laperche Y et al. Gas6 deficiency prevents liver inflammation, steatohepatitis, and fibrosis in mice. Am J Physiol Gastrointest Liver Physiol 2011; 300: G1043–1053.

32 Korshunov VA, Mohan AM, Georger MA, Berk BC. Axl, a receptor tyrosine kinase, mediates flow-induced vascular remodeling. Circ Res 2006; 98: 1446–1452.

33 Melaragno MG, Fridell YW, Berk BC. The Gas6/Axl system: a novel regulator of vascular cell function. Trends Cardiovasc Med 1999; 9: 250–253.

34 O’Donnell K, Harkes IC, Dougherty L, Wicks IP. Expression of receptor tyrosine kinase Axl and its ligand Gas6 in rheumatoid arthritis: evidence for a novel endothelial cell survival pathway. Am J Pathol 1999; 154: 1171–1180.

35 Antony J, Tan TZ, Kelly Z, Low J, Choolani M, Recchi C et al. The GAS6-AXL signaling network is a mesenchymal (Mes) molecular subtype-specific therapeutic target for ovarian cancer. Sci Signal 2016; 9: ra97.

36 Wilson C, Ye X, Pham T, Lin E, Chan S, McNamara E et al. AXL inhibition sensitizes mesenchymal cancer cells to antimitotic drugs. Cancer Res 2014; 74: 5878–5890.

37 Diepenbruck M, Christofori G. Epithelial-mesenchymal transition (EMT) and metastasis: yes, no, maybe? Curr Opin Cell Biol 2016; 43: 7–13.

38 Saitoh M. Involvement of partial EMT in cancer progression. J Biochem 2018; 164: 257–264.

39 Kim YS, Jung SH, Jung DH, Choi SJ, Lee YR, Kim JS. Gas6 stimulates angiogenesis of human retinal endothelial cells and of zebrafish embryos via ERK1/2 signaling. PLoS One 2014; 9: e83901.

40 Zuo PY, Chen XL, Lei YH, Liu CY, Liu YW. Growth arrest-specific gene 6 protein promotes the proliferation and migration of endothelial progenitor cells through the PI3K/AKT signaling pathway. Int J Mol Med 2014; 34: 299–306.

41 Stenhoff J, Dahlback B, Hafizi S. Vitamin K-dependent Gas6 activates ERK kinase and stimulates growth of cardiac fibroblasts. Biochem Biophys Res Commun 2004; 319: 871–878.

42 Leake I. Pancreatic cancer: surprising role for fibrosis. Nat Rev Gastroenterol Hepatol 2014; 11: 396.

43 Ozdemir BC, Pentcheva-Hoang T, Carstens JL, Zheng X, Wu CC, Simpson TR et al. Depletion of Carcinoma-Associated Fibroblasts and Fibrosis Induces Immunosuppression and Accelerates Pancreas Cancer with Reduced Survival. Cancer Cell 2015; 28: 831–833.

44 Rhim AD, Oberstein PE, Thomas DH, Mirek ET, Palermo CF, Sastra SA et al. Stromal elements act to restrain, rather than support, pancreatic ductal adenocarcinoma. Cancer Cell 2014; 25: 735–747.

45 Weniger M, Honselmann KC, Liss AS. The Extracellular Matrix and Pancreatic Cancer: A Complex Relationship. Cancers (Basel) 2018; 10.

46 Cabezon R, Carrera-Silva EA, Florez-Grau G, Errasti AE, Calderon-Gomez E, Lozano JJ et al. MERTK as negative regulator of human T cell activation. J Leukoc Biol 2015; 97: 751–760.

47 Walzer T, Vivier E. NK cell development: gas matters. Nat Immunol 2006; 7: 702–704.

48 Feig C, Gopinathan A, Neesse A, Chan DS, Cook N, Tuveson DA. The pancreas cancer microenvironment. Clin Cancer Res 2012; 18: 4266–4276.

49 Hingorani SR, Wang LF, Multani AS, Combs C, Deramaudt TB, Hruban RH et al. Trp53(R172H) and KraS(G12D) cooperate to promote chromosomal instability and widely metastatic pancreatic ductal adenocarcinoma in mice. Cancer Cell 2005; 7: 469–483.

50 Finck R, Simonds EF, Jager A, Krishnaswamy S, Sachs K, Fantl W et al. Normalization of mass cytometry data with bead standards. Cytometry Part A 2013; 83A: 483–+.

51 Schmittgen TD, Livak KJ. Analyzing real-time PCR data by the comparative C-T method. Nature Protocols 2008; 3: 1101–1108.

