## Supplementary figures and images for "Blockade of stromal Gas6 alters cancer cell plasticity, activates NK cells and inhibits pancreatic cancer metastasis"

### Supplemental Figure 1

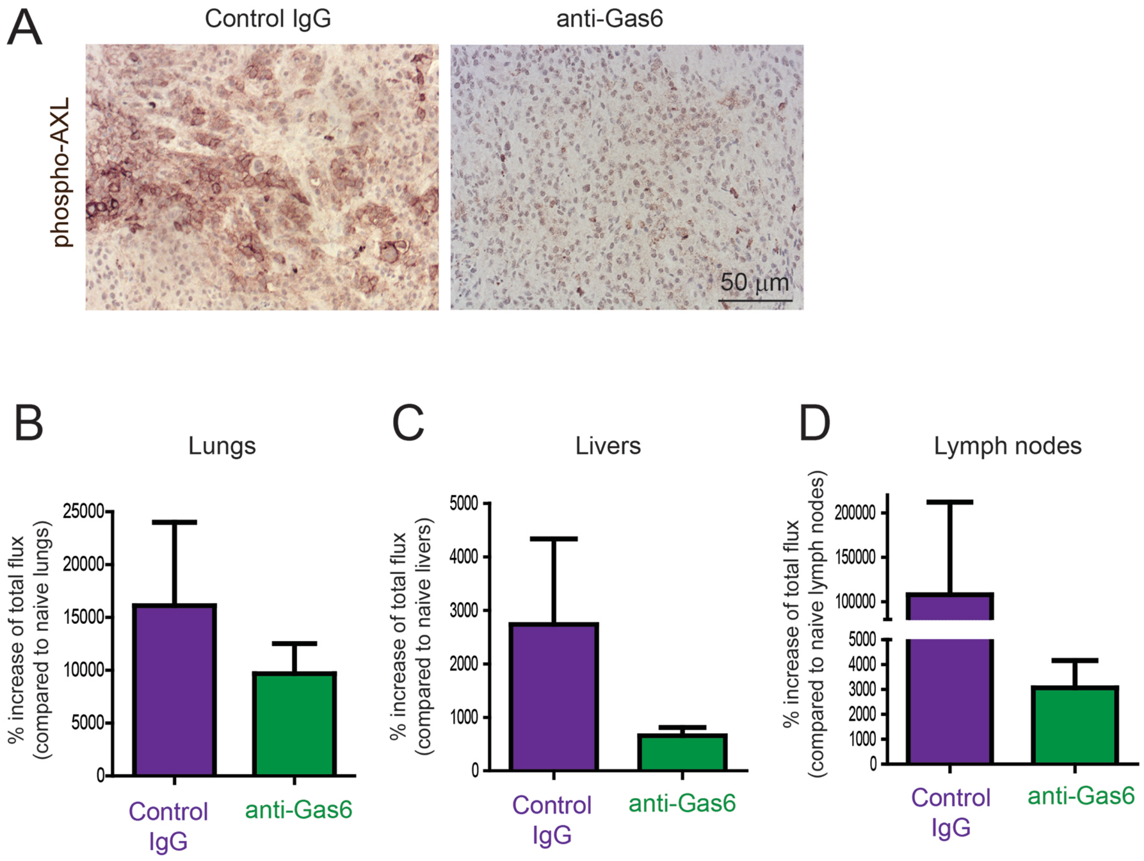

### Supplemental Figure 2

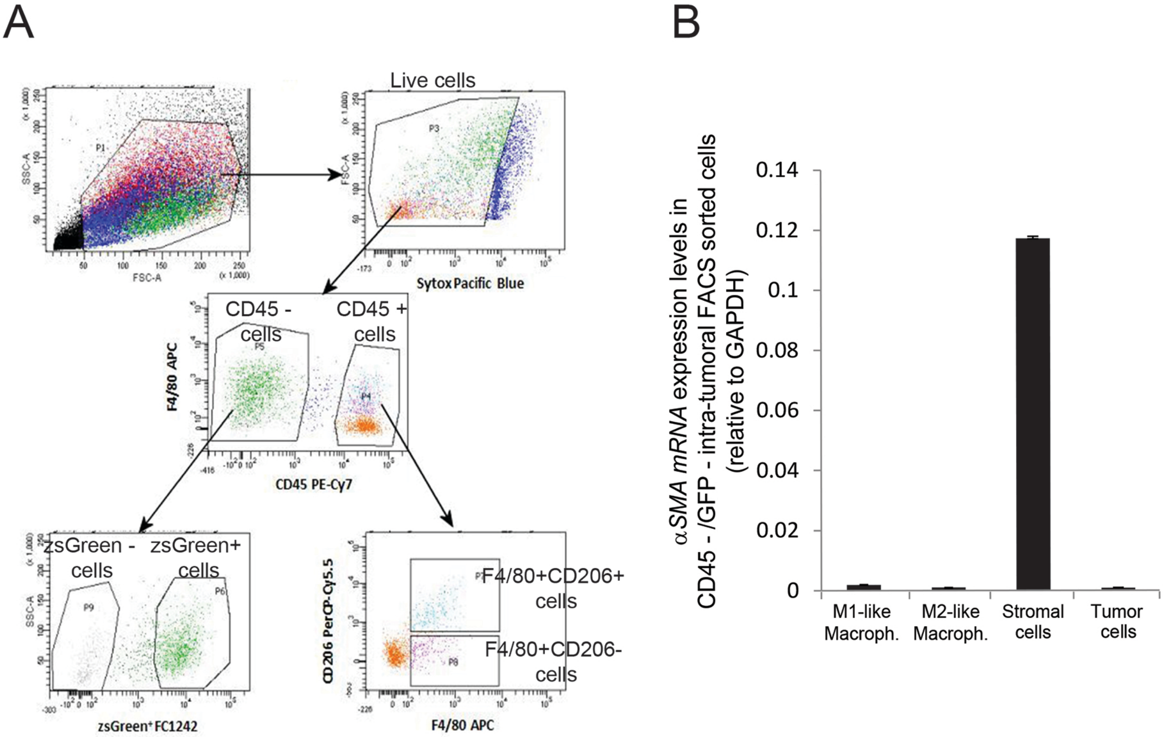

### Supplemental Figure 4

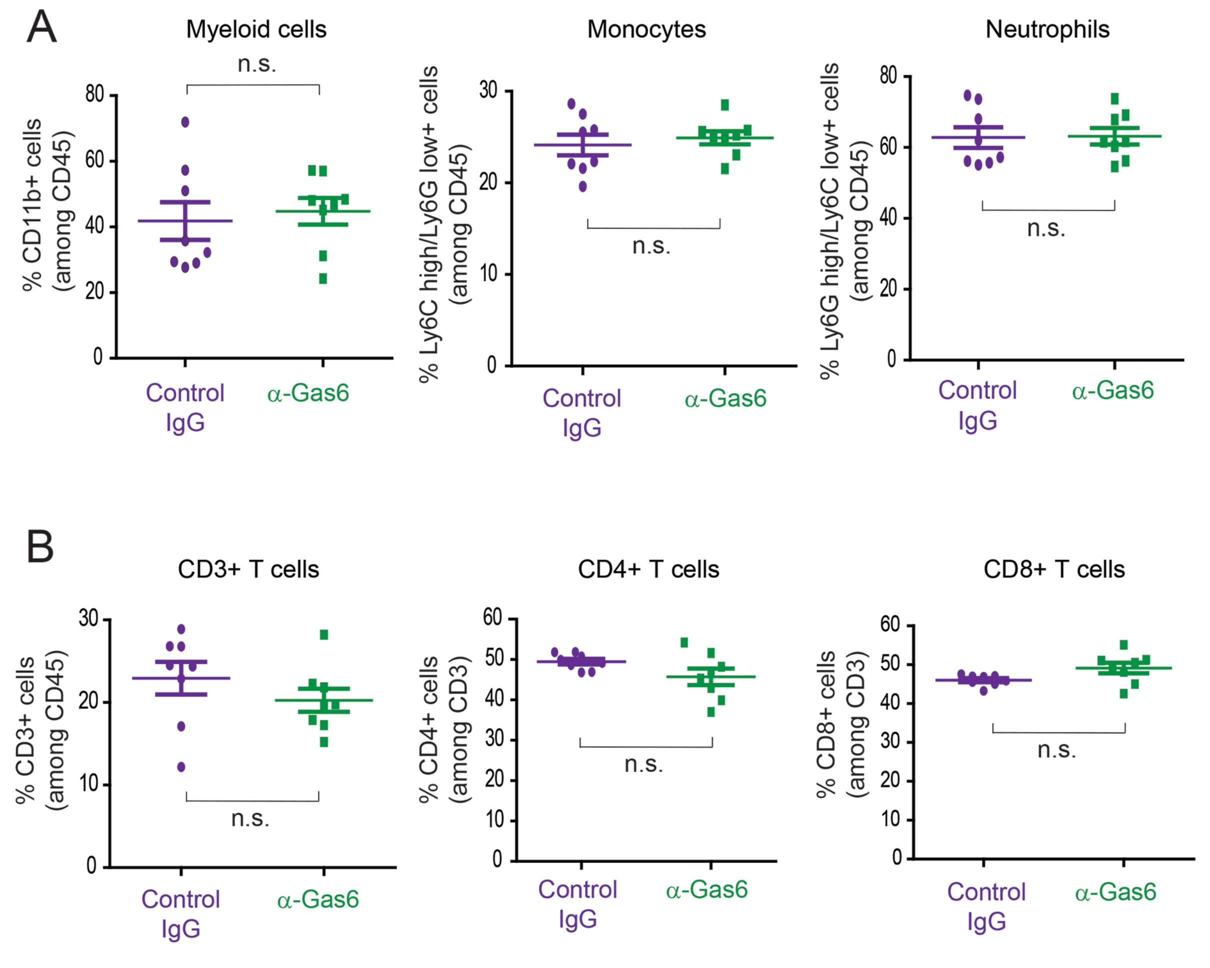

### Supplemental Figure 5

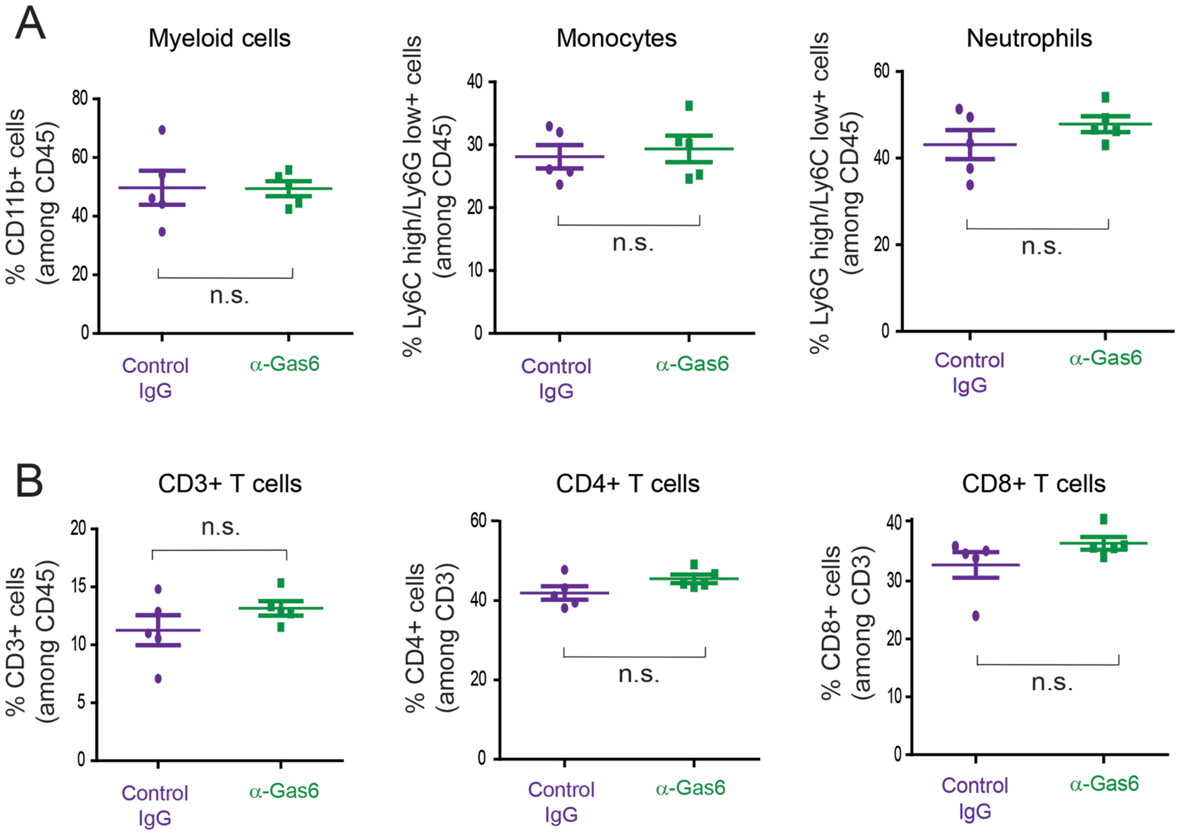

### Supplemental Figure 6

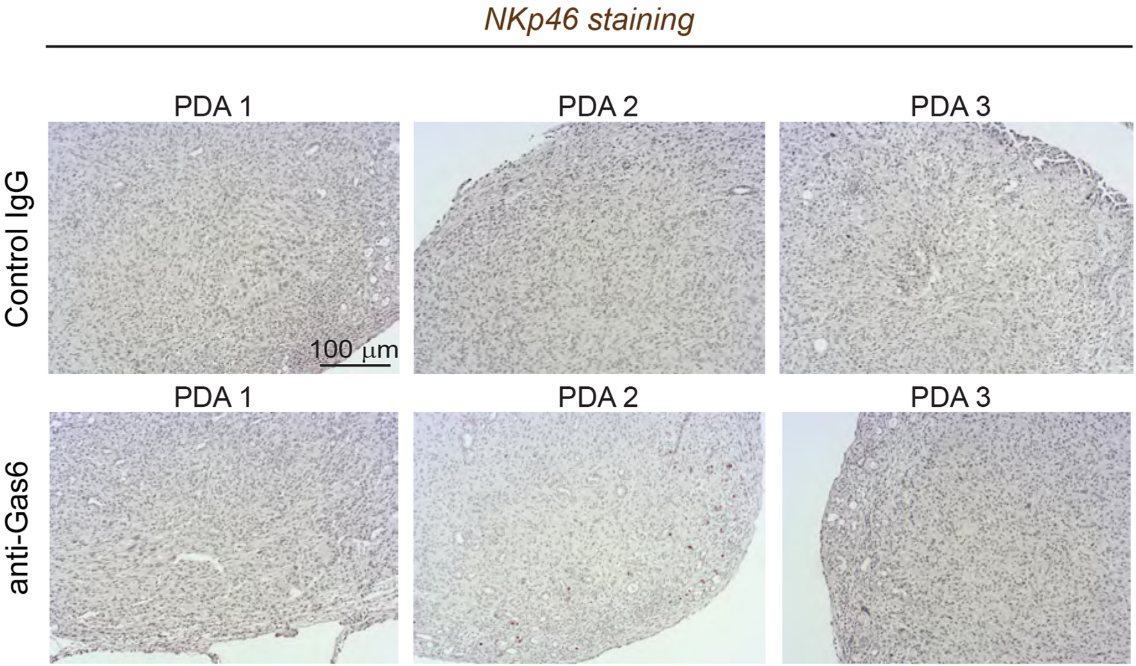
