## Supplemental Figure 3 for "Blockade of stromal Gas6 alters cancer cell plasticity, activates NK cells and inhibits pancreatic cancer metastasis"

A

| Cell Type | Marker | Clone | Tag |
|---|---|---|---|
| **Immune cells** | CD45 | 30-F11 | 147 Sm |
| **T cells** | CD3e | 145-2C11 | 152 Sm |
| | CD4 | RM4-5 | 145 Nd |
| | CD8a | 53-6.7 | 168 Er |
| | CD25 (IL-2R) | 3C7 | 151 Eu |
| | CD44 | IM7 | 171 Yb |
| | CD62L (L-selectin) | MEL-14 | 160 Gd |
| **T cell activation/exhaustion** | CD69 | H1.2F3 | 143 Nd |
| | CD279 (PD-1) | RMP1-30 | 159 Tb |
| | CD152 (CTLA-4) | UC10-4B9 | 154 Sm |
| | CD274 (PD-L1) | 10F.9G2 | 153 Eu |
| **Myeloid cells** | I-A/I-E (MHC class II) | M5/114.15.2 | 174 Yb |
| | CD11b (Mac-1) | M1/70 | 148 Nd |
| | F4/80 | BM8 | 146 Nd |
| | CD86 | GL1 | 172 Yb |
| | CD206 | C068C2 | 169Tm |
| | CD11c | N418 | 142 Nd |
| | CD11b (Mac-1) | M1/70 | 148 Nd |
| | Ly-6G | 1A8 | 141 Pr |
| | Ly-6C | HK1.4 | 162 Dy |
| | CD115 | AFS98 | 144 Nd |

B

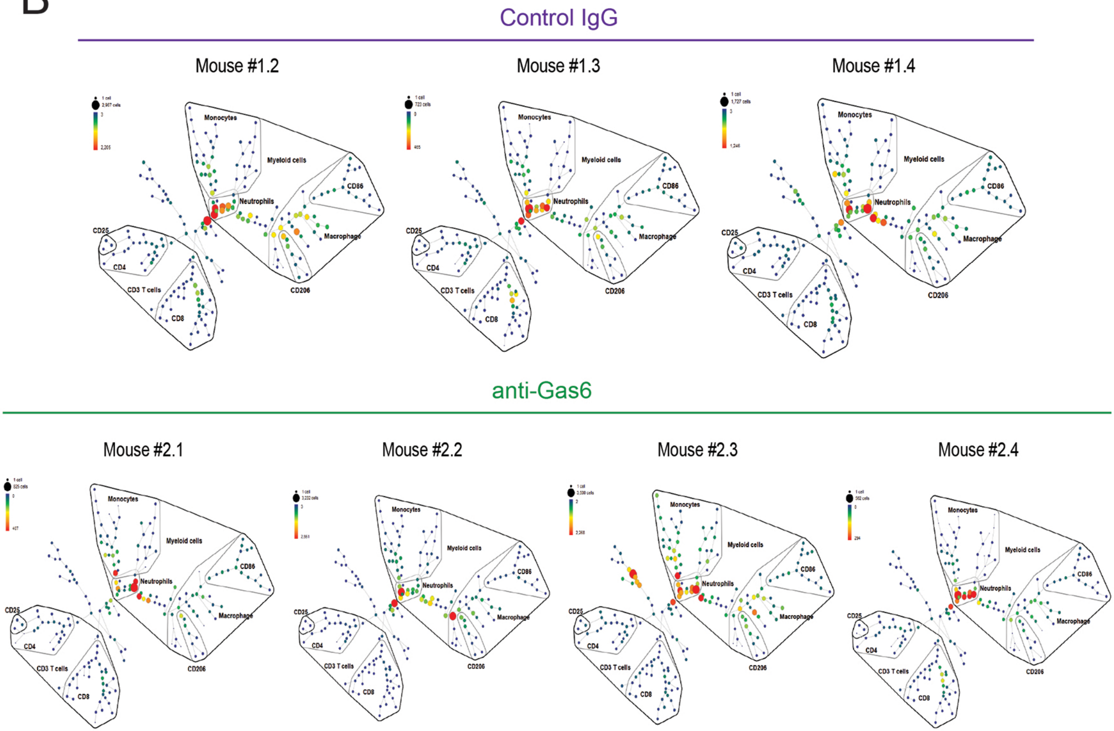
