## Supplemental Figure Legends for "Blockade of stromal Gas6 alters cancer cell plasticity, activates NK cells and inhibits pancreatic cancer metastasis"

### Supplementary Figure legends

**Supplementary Figure 1. Anti-Gas6 treatment reduces metastasis.**

**(A)** Immunohistochemical staining of phospho-AXL in pancreatic tumors treated with IgG (control) or anti-Gas6 antibody. Scale bars 100 μm and 50 μm**. (B)** Quantification of metastasis in lungs, **(C)** livers and **(D)** mesenteric lymph nodes by *ex-vivo* bioluminescent signalling (IVIS imaging technology).

**Supplementary Figure 2. Gating strategy used to FACS-sort tumor cells, tumor associated macrophages and stromal cells from PDA tumors.**

**(A)** FACS gating strategy of PDA tumours for tumor cells (Sytox-, CD45-, zsGreen+), non-immune stromal cells (Sytox-, CD45-, zsGreen-), M1-like macrophages (Sytox-, CD45+, F4/80+, CD206-) and M2-like macrophages (Sytox-, CD45+, F4/80+, CD206+). **(B)** qPCR analysis of *αSMA* expression relative to GAPDH in FACS sorted M1-like and M2-like macrophages, stromal cells and tumor cells. Values shown are the mean and SEM (n=3).

**Supplementary Figure 3. SPADE analysis of CyTOF analysed PDA tumors.**

**(A)** Table of heavy metal-conjugated antibodies used for CyTOF analysis, all antibodies were purchased from Fluidigm. **(B)** Control IgG treated and anti-Gas6 treated mice SPADE tree figures of CD45+ cells. Manual gating identified myeloid cells (MHC II+, CD11b+) monocytes (Ly6C high/Ly6G low), neutrophils (Ly6C low/Ly6G high), M1-like macrophages (F4/80+, CD68+), M2-like macrophages

(F4/80+, CD206+) and T cells (CD3+): T helper (CD4+), Cytolytic T cells (CD8+) and T regulatory cells (CD25+).

**Supplementary Figure 4. Anti-Gas6 treatment does not alter myeloid cell or T cell levels in the peripheral blood**

**(A)** FACS analysis of Control IgG and anti-Gas6 treated mice peripheral blood (n=8 per treatment group) for the presence of myeloid cells (CD11b+), monocytes (Ly6C high/Ly6G low) and neutrophils (Ly6C low/Ly6G high). **(B)** T cell populations: pan-T cell (CD3+), T regulatory cells (CD4+) and cytotoxic T cells (CD8+).

**Supplementary Figure 5. Anti-Gas6 treatment does not alter myeloid cell or T cell levels in the lungs**

**(A)** FACS analysis of Control IgG and anti-Gas6 treated mice lung metastasis (n=5 per treatment group) for presence of myeloid cells (CD11b+), monocytes (Ly6C high/Ly6G low+) and neutrophils (Ly6C low/Ly6G high+). **(B)** T cell populations: pan-T cell (CD3+), T regulatory cells (CD4+) and cytotoxic T cells (CD8+).

**Supplementary Figure 6. Anti-Gas6 treatment does not increase NK cell numbers in primary pancreatic tumors.**

Representative immunohistochemical staining of NK cells in primary pancreatic tumors from mice treated with control IgG or anti-Gas6 antibody. Scale bar, 100 μm.
